# Conserved SQ and QS motifs in bacterial effectors suggest pathogen interplay with the ATM kinase family during infection

**DOI:** 10.1101/364117

**Authors:** Davide Sampietro, Hugo Sámano-Sánchez, Norman E. Davey, Malvika Sharan, Bálint Mészáros, Toby J. Gibson, Manjeet Kumar

**Affiliations:** Department of Biotechnology and Biosciences, University of Milano-Bicocca, Piazza della Scienza 2, 20126 Milan, Italy.; Structural and Computational Biology Unit, European Molecular Biology Laboratory, Heidelberg, 69117, Germany; (Candidate for) Joint PhD degree from EMBL and Heidelberg University, Faculty of Biosciences.; UCD School of Medicine & Medical Science, University College Dublin, Belfield, Dublin 4, Ireland.; MTA-ELTE Momentum Bioinformatics Research Group, Department of Biochemistry, Eötvös Loránd University, Budapest H-1117, Hungary.

## Abstract

Understanding how bacteria hijack eukaryotic cells during infection is vital to develop better strategies to counter the pathologies that they cause. ATM kinase family members phosphorylate eukaryotic protein substrates on Ser or Thr residues followed by Gln. The kinases are active under oxidative stress conditions and/or the presence of ds-DNA breaks. While examining the protein sequences of well-known bacterial effector proteins such as CagA and Tir, we noticed that they often show conserved (S/TQ) motifs, even though the evidence for effector phosphorylation by ATM has not been reported. We undertook a bioinformatics analysis to examine effectors for their potential to mimic the eukaryotic substrates of the ATM kinase. The candidates we found could interfere with the host’s intracellular signaling network upon interaction, which might give an advantage to the pathogen inside the host. Further, the putative phosphorylation sites should be accessible, conserved across species and, in the vicinity to the phosphorylation sites, positively charged residues should be depleted. We also noticed that the reverse motif (QT/S) is often also conserved and located close to (S/TQ) sites, indicating its potential biological role in ATM kinase function. Our findings could suggest a mechanism of infection whereby many pathogens inactivate/modulate the host ATM signaling pathway.

## INTRODUCTION

Clinical data have uncovered a clear connection between several molecular-level biological processes in the human cell, such as oxidative stress response and DNA damage repair, and pathogenic infections. Specifically, the exact molecular steps through which pathogens modulate such host processes are largely unknown. However, several human kinases are known to be critical components of affected pathways and they are considered as candidate targets for pathogenic intervention.

Phosphatidylinositol-3 kinase-related kinases (PIKKs) encompass several Ser/Thr-protein kinases which share sequence similarity with lipid phosphorylating phosphatidylinositol-3 kinases (PI3Ks) (1). In humans, there are six different PIKK members: Ataxia-Telangiectasia Mutated (ATM), Ataxia- and Rad3-related (ATR), DNA-dependent Protein Kinase catalytic subunit (DNA-PK), mammalian Target Of Rapamycin (mTOR), Suppressor of Morphogenesis in Genitalia (SMG-1) and Transformation/Transcription domain-Associated Protein (TRRAP) (1). ATM, ATR and DNA-PK are involved both in the cell cycle DNA damage checkpoints (2, 3) and more generally in oxidative stress response (4; 5). Even though ATM is mainly considered a nuclear cell cycle kinase, there are several pieces of evidence suggesting it is also localised and functional in the cytosol (6 - 8). Similarly, there is experimental evidence for ATR also being present in the cytosol, although its role is still poorly understood (9). Additionally, DNA-PK was reported to be present in the cytosol where it can activate the transcription of type I interferon (IFN), cytokine and chemokine genes during viral infections (10).

Patients with ATM mutations suffering from Ataxia Telangiectasia (AT) display a progressive loss of neurons (11). Given that neurons are non-dividing cells, this AT phenotype is an indication that ATM has roles outside of cell cycle checkpoints. AT patients are known to be immunodeficient and are especially susceptible to pulmonary infections (12,13). Airway epithelial cells of AT patients have been found to be very sensitive to oxidative insult during *Streptococcus pneumoniae* infection (14). Certain pathogens deliberately exploit the DNA damage signaling systems. Several of them, including certain *Escherichia coli* strains, inject genotoxins into host cells, such as the Cytolethal Distending Toxin (CDT) nucleases which create double-strand DNA (ds-DNA) breaks: the resultant ATM damage response can lead to apoptosis of the targeted cells (15). By contrast, Human papillomavirus 16 (HPV16) requires a more controlled ATM activation of the DNA damage repair system for viral genome activation (16).

ATM, ATR and DNA-PK (hereafter referred to as AADks) phosphorylate a wide range of substrates at a Ser-Gln or Thr-Gln (pS/TQ) motif (17). However, in addition to this minimal essential dipeptide, there are preferences for hydrophobic and negatively charged residues N-terminal to the S/TQ site; conversely, positively charged residues are underrepresented in the proximity of the phosphorylation (S/TQ) site (17). SQ and TQ are examples of Short Linear Motifs (SLiMs), which are small modules of a protein sequence mediating protein-protein interactions. SLiMs are frequently located in intrinsically disordered protein (IDP) regions. Since disordered regions are not involved in stably folded protein structure formation, mutations are usually more tolerated: Residue conservation in these regions is expected to be indicative of function (18,19). Several known substrates of AADks have clusters of many S/TQs, for example BRCA1, Chk2 and most strikingly FUS (20). However, it is currently unclear what is the cellular purpose for those clusters of SQ sites, given that other AADks substrates, such as H2AX and p53, can fulfil their biological roles by having just a single major site (21).

Proteins such as CDT that are introduced into the host cell by bacteria via various secretory systems are termed effectors (22). Using effectors, pathogenic bacteria can hijack the host cell system in order to sustain themselves against the immune response and favour their own replication in a series of processes called host-pathogen interactions (23). Up to now, six different secretory systems (TSS) have been described in Gram-negative bacteria (24); Gram-positive bacteria can use either a specialised apparatus called the injectisome or use additional components in order to introduce the effector into the cytoplasm of the host cell (25). Apart from the six TSS’s in Gram-negative bacteria, a seventh one has also been identified exclusively in the members of the genus Mycobacteria (26). In addition, bacteria may use other ways to secrete proteins such as the classical secretion pathway (Sec) or the twin-arginine pathways (Tat pathways) (27,28). Despite our extensive knowledge of TSS, for many effectors the exact mechanism of secretion is still unknown. T3SS, T4SS and T6SS can cross both the bacterial and eukaryotic cellular membrane (24). Even though translocating a protein is energetically expensive (29), *Enteropathogenic E. coli* (EPEC) uses T3SS to translocate the Translocated-intimin receptor (Tir) effector into the host plasma membrane of intestinal epithelial cells. Upon secretion, it interacts with cytoplasm-located TNF α receptor-associated factor (TRAF) causing its degradation and the inactivation of NF-κB (30). Other bacteria use T4SS: this is the case for *Helicobacter pylori*, which employs this system to introduce CagA into gastric cells, thereby making it able to interact with a set of host cell regulatory proteins (31). Further, as with viruses (32), bacterial effector proteins can also mimic SLiMs in order to successfully carry out infection. The concept of molecular mimicry is well established for CagA (33,34) and Tir (35). We have been working to collate these, along with other known effector SLiM mimics into the ELM resource (36).

While reviewing the effectors with published SLiMs, we noticed that many of them, including CagA and Tir had multiple conserved or semi-conserved S/TQs (Table S1). We also noticed that they had conserved or semi-conserved QS/T motifs, having no associated function yet. When we checked cellular FUS and other ATM substrates, we found that the QS/T motif was also present in multiple copies, suggesting it might have a functional relationship with S/TQs. In this paper, we report a bioinformatics analysis of these motifs in cellular proteins and effectors. Because these two residue motifs are so simple and Ser is the most frequent amino acid found in proteins, there is very little sequence information content to work with, and it is not straightforward to obtain good statistical support. Nevertheless, it is clear that a very broad property of invasive bacterial pathogens is that they have effectors with S/TQ and QS/T motifs, and therefore they may be interacting with the AADks to mimic cellular substrates. Experimental investigation on whether or not these motifs are phosphorylated is desirable, as it might open up new therapeutic strategies.

## MATERIALS AND METHODS

Experimentally validated phosphorylation sites in the substrates of ATM, ATR and DNA-PK were retrieved from PhosphositePlus (37) and Phospho.ELM (38). Mouse and rat sequences, human alternative isoforms and autophosphorylation sites were removed. In total, we collected 152 proteins containing 360 experimentally validated pS/TQ sites (Supplementary dataset 1).

The experimentally validated effectors for T3SS, T4SS and T6SS were collected from SecretEPDB (39). Low quality (such as partial sequences) and redundant sequences were removed at 90% sequence identity cut-off. After the filters, we gathered 1,000 unique sequences (Supplementary dataset 2).

Protein sequences were collected from the UniProt database (40) and were aligned with Clustal Omega (41). The alignments were visualised using the Jalview software (42). To measure sequence conservation, the probability of relative local conservation (43) was used. To create the negative control for the S/TQ and QS/T site conservation, the entire Human proteome (SwissProt) was downloaded. Subsequently, we extracted proteins having the following gene ontology accession - GO:0005737 (cytoplasm), GO:0005829 (cytosol), GO:0005634 (nucleus), and GO:0005622 (intracellular). From this subset, we removed all the 152 experimentally validated substrates of AADks and all the AADk interactors from IntAct and BioGRID. We collected 8,700 proteins that are not known to interact directly with AADks.

The significance of each peptide was calculated as the distance from a 1:1 correlation line as follows:

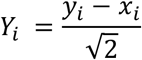

Where Y_i_ is the distance from the 1:1 correlation line of the peptide i, x_i_ and y_i_ are the frequencies of the peptide i in the scrambled and original sequences. A Y_i_ higher than 0 means that the peptide i is overrepresented in the positive control; on the contrary, a lower Y_i_ than 0 indicates that i is enriched in the negative control.

In order to detect the disordered regions of a protein, we used the IUPred software (44). Two minimal lengths (8 and 15 amino acids) for the disordered regions were tested. The frequencies of all the 400 possible dipeptides in the disordered region of the human proteome did not vary substantially (Pearson’s correlation for the minimal window of 8 versus a minimal window of 15 was > 0.99). Therefore, a disordered region was defined as a sequence with at least 15 consecutive amino acids with an IUPred score ≥ 0.4. In the mentioned window up to two residues with scores below the threshold were allowed. Statistical analyses were run in R and Python (Numpy and Scipy libraries).

## RESULTS

The increasing use of infectious tissue culture systems has been helpful in identifying bacterial effectors. For some of these, linear motif mimicry has been reported, and so we have been collecting these motifs to enter into the ELM database. One example is the EPIYA motif which occurs as repeats in CagA, where it was found to mimic an eukaryotic substrate (45). While studying conservation of this motif to prepare the ELM resource entry, we noticed that CagA alignments also had conserved SQs, despite a lack of evidence for ATM phosphorylation (Figure S1). We continued to see SQs in other effectors. An example of a bacterial effector, lpg2844, with a cluster of S/TQs is shown in Figure 1A. It was also evident that the (reverse) QS/T motif also repeats and clusters with the S/TQs (Figure 1). We then realised that the S/TQs and Q/STs cluster in some cellular ATM substrates as well. The cellular protein FUS has twelve S/TQ motifs clustered in the first third quarter of the protein sequence together with a similar number of QS/Ts (Figure 1B). Other effectors, including PopA, PopB, Tir and yhhA, are shown in Figure 1C, highlighting the clusters of S/TQ and QS/T in their sequences. Although there is no evidence for functional equivalence, these sequence similarities are so striking that they raised questions about whether bacterial effector proteins are generally rich in S/TQ motifs and whether the QS/T motif might be functional in cellular AADk substrates. The answer to these questions will require experimental testing. Therefore, we were interested in undertaking a closer look bioinformatically to see if there is statistical support for these motif associations that would warrant the subsequent experimental work.

**Figure 1.**
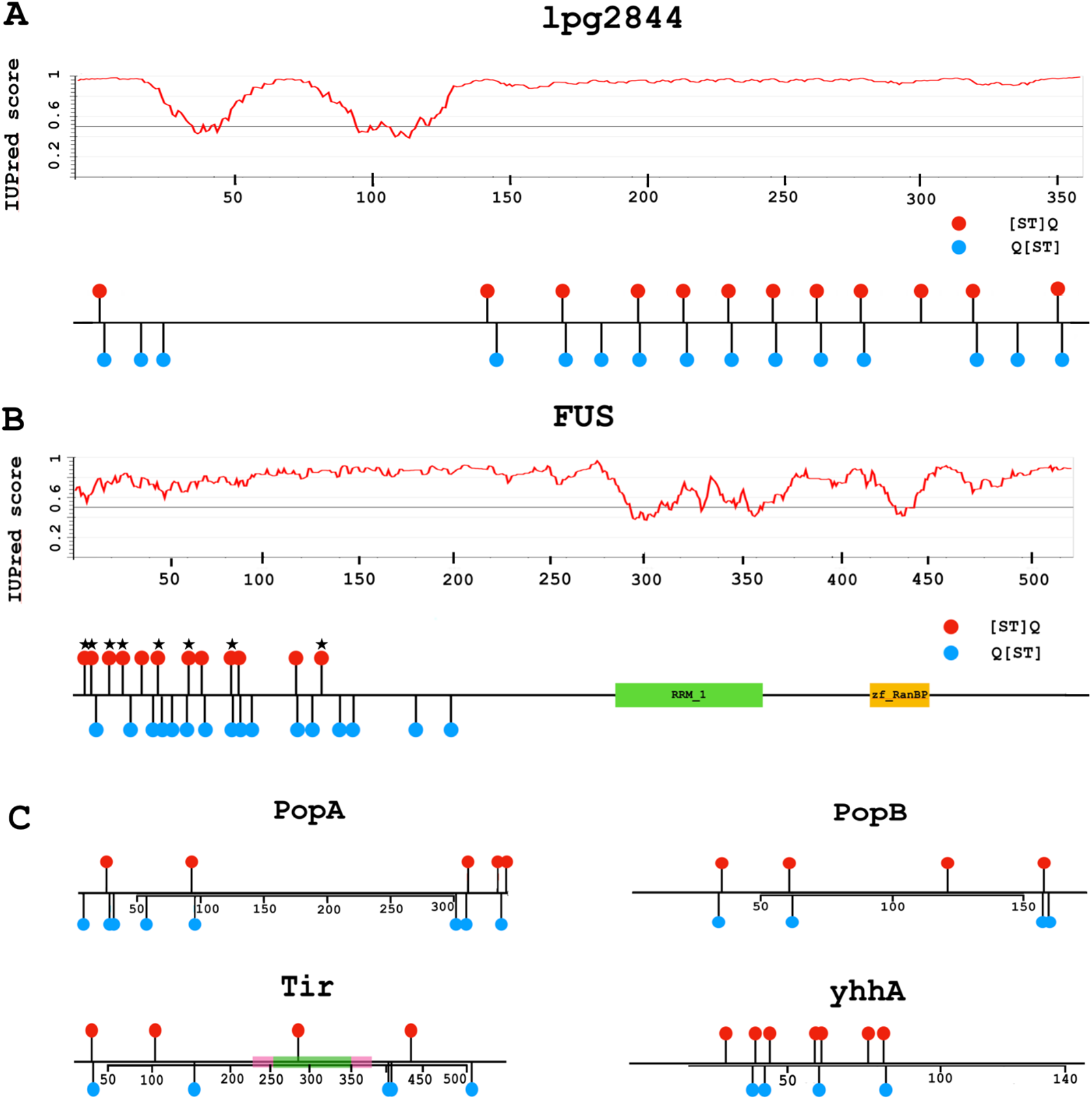
Clustering of S/TQ (red lollipop) and QS/T (blue lollipop) motifs in bacterial effectors and in a cellular ATM substrate. The IUPred plot shows that in both proteins, the motifs are in regions of strong predicted native disorder (score well above 0.5). **A:** The bacterial effector lgp2844 (UniProt:Q5ZRN6) from *Legionella pneumophila* is a potential but unproven candidate for a function with FUS-like motif mimicry. **B:** The subset of S/TQ motifs in the cellular ATM substrate FUS (UniProt:P35637) which are known to be phosphorylated by AADks are marked with a star. The function of the remaining motifs is unknown. **C:** Other interesting examples include PopA (Q9RBS0) and PopB (Q9RBS1) From *Ralstonia solanacearum*, Tir (B7UM99) and yhhA (B7UKZ9) from *E.coli*. For Tir, the topological domains are shown (green for extracellular, pink for helical trans-membrane). It should be noted that the SQ in the extracellular domain of Tir cannot be phosphorylated by AADks.

### SQs are enriched in the disordered regions of the substrates

Very short motifs, such as SQ and QS have low information content. This makes the reliable assessment of the significance of their occurrence challenging. Known functional SQ motifs reside in disordered regions, and considering disordered protein segments in general, Ser and Gln are known to be disorder-promoting (46). A comparison between the disordered regions of the known cellular AADk substrates and the disordered regions of the whole human proteome revealed that Ser, Gln and Thr are the three most overrepresented residues in the substrates (Figure 2). This raises the question whether the high observed number of S/TQ sites is significant or is it purely by random chance in the disordered regions of the AADk substrates. For this, the number of occurrences in the real sequences was compared to occurrences in a set of background sequences, generated by shuffling the sequences of the disordered regions of each substrate 10,000 times (scrambled sequences). By scrambling the sequences, any resultant S/TQs have appeared by chance. The occurrences in these randomized sequences model the theoretical frequencies of SQs and QSs (together with the frequencies of all 400 possible dipeptides). We found that SQ sites are enriched in the substrates (1 - ecdf(SQ) = 0.0225) (Figure 3A, Figure S2); however, QS and QT motifs are underrepresented compared to randomized sequences.

**Figure 2.**
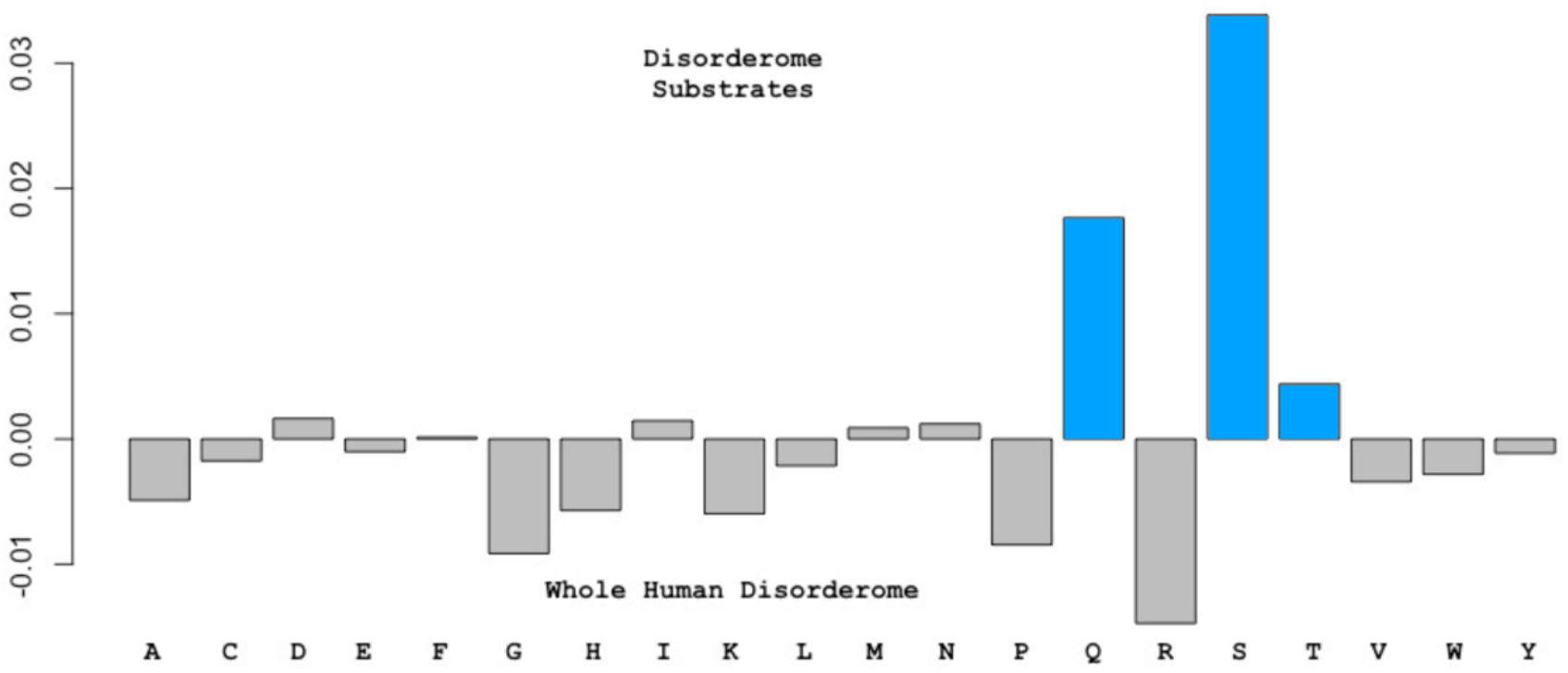
Amino acid frequency in the disordered regions of the substrates versus the whole human disorderome. Each bar represents one amino acid. The relative frequencies are shown in the y-axis and a value higher than 0 means that the residue is overrepresented in the disordered regions of the substrates. It should be noted that S, Q and T (light blue bars) are the most abundant residues.

**Figure 3.**
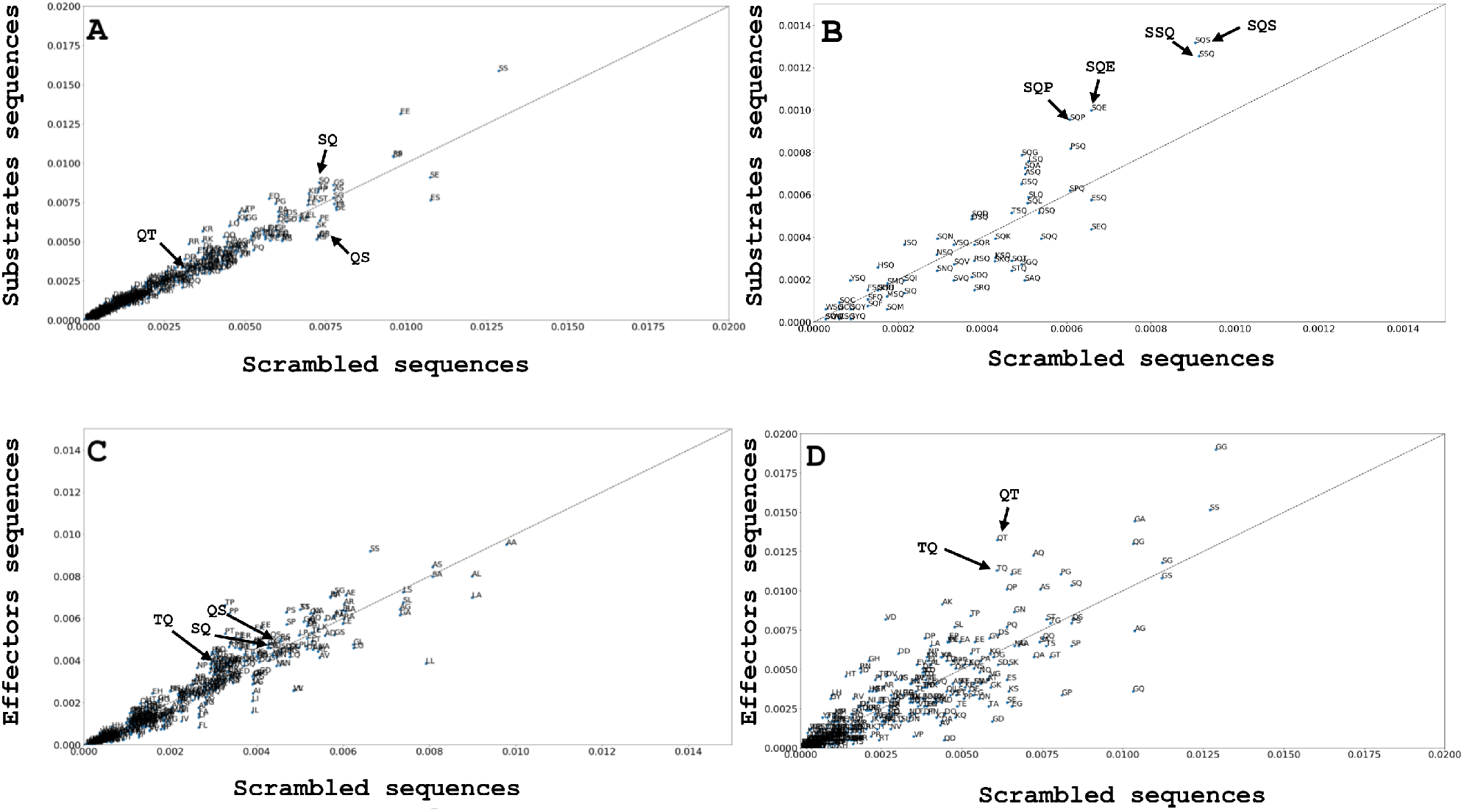
A and B show the frequency of all dipeptides and tripeptides matching the SxQ, SQx and xSQ regular expression (where x corresponds to any amino acid) in the AADk substrates. C displays the frequency of each dipeptide in the entire set of effectors, whereas D in our effector candidates. The diagonal is the 1:1 correlation line. The further the dot is from this diagonal, the stronger the enrichment. Whatever is above the diagonal is enriched in the positive set.

The consideration of tripeptides provides an auxiliary test for the biological relevance of SQ and QS occurrences. The high fraction of S and Q residues are expected to yield a high number of randomly occurring SxQ tripeptides (x indicates any amino acid) without biological meaning. In contrast, SQx or xSQ tripeptides trivially contain all functional SQ sites. Comparing the occurrence of tripeptides, we found that SxQ sites are less common than SQx and xSQ, reflecting the abundance of SQ motifs with biological roles (Figure 3B, Figure S3). In particular, SQS, SQP, SQE, and SSQ are the top-scoring tripeptides (1 - ecdf(SQS, SQP, SQE, SSQ) = <0.0166, 0.0166, 0.033, 0.5, respectively) (Figure 3B). Similarly, considering the occurrences of tetrapeptides (e.g. SxxQ), the top-scoring peptides corresponded to the placement of x’s that do not destroy the SQ motif (Figure S4). Altogether, these data show that the over-representation of S/TQs in the AADk substrates is not simply a consequence of the high proportion of Ser and Gln residues in the disordered regions, but reflects true biological function. While checking the frequency of occurrence of the S/TQ dipeptides in effectors, we observed that not all the effectors followed the same distribution pattern as seen in AADk substrates (Figure 3C). This can be possible as it seems unlikely that all effectors should act to mimic the substrates of AADks (Figure 3D). Still, we noticed that in some bacterial effectors the TQ and QT dipeptides also occur more frequently than in the scrambled set (1-ecdf(QT) < 0.0025, 1-ecdf(TQ) = 0.0075).

### Uneven distribution of SQ-QS distances indicates a possible biological role

Our observation that S/TQs often appear close to QS/Ts in the sequences of the substrates and the effectors (Figure 1, Figure 4, Figure S5-S7) motivated us to check the distances between S/TQs and QS/Ts in general. The distance between an S/TQ site and the closest QS/T site is hereafter referred to as “SQ-QS distance”. The significance assessment of SQ-QS distances was performed by calculating the p-values using the Chi-square test. The theoretical distribution of the SQ-QS distances was compared to both the distribution of SQ-QS distances from the experimentally validated phosphosites and the distribution of the SQ-QS distances from all the ST/Qs and QS/Ts in the disordered regions of the substrates. We observed that the p-value was lower in the former than in the latter (Chi-square test p-value = 0.00631 and 0.0278 respectively; Figure S8). The lower p-value of SQ-QS distances in the experimentally validated phosphosites suggests a previously unknown potential biological function for QS/T, that is correlated with the function of S/TQ sites.

**Figure 4.**
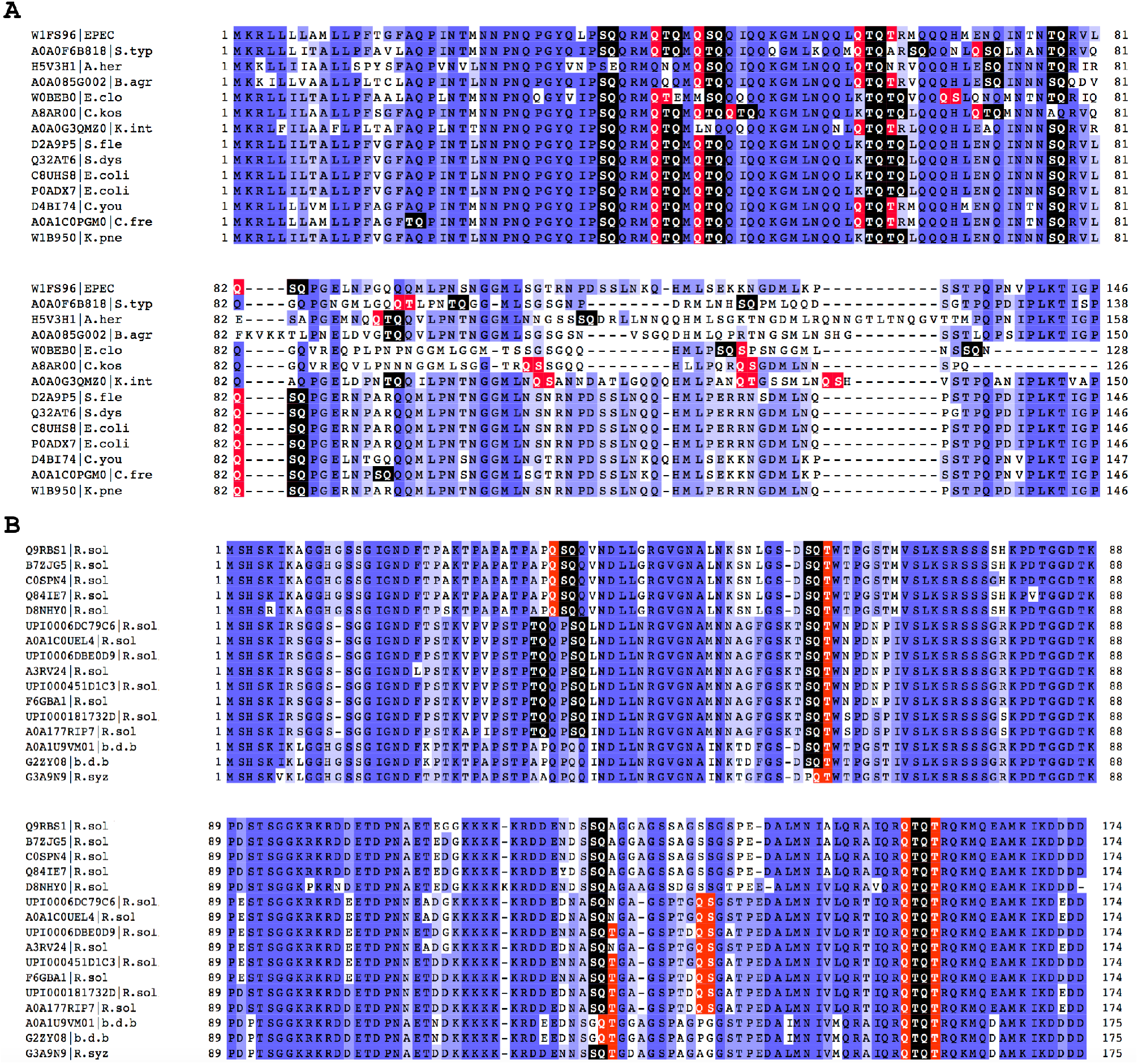
Panel A shows an alignment for yhhA from *Enteropathogenic E.coli* (EPEC), and its homolog from *Atlantibacter hermanni* (A.her), *Buttiauxella agrestis* (B.agr), *Enterobacter cloacae* (E.clo), *Citrobacter koseri* (C.kos), *Kluyvera intermedia* (K.int), *Shigella flexneri* (S.fle), *Shigella dysenteriae* serotype 1 (S.dys), *Escherichia coli* (E.coli), *Salmonella enterica* Typhimurium (S.typ), *Citrobacter freundii* (C.fre), *Klebsiella pneumoniae* (K.pne). Panel B shows possible AADk phosphorylation sites in PopB (Q9RBS1) from different strains of *Ralstonia solanacearum* (R.sol), *Banana Blood Disease Bactetrium* (b.d.b) and *Ralstonia syzygii* (R.syz). S/ TQs are highlighted in black, QS/Ts in red. The motifs cluster together and often overlap. These proteins are predicted to be natively disordered. Blue marks the percentage of identity for all other residues. Uniprot and UniRef accessions have been provided for the sequences.

### pS/TQ sites are embedded in characteristic protein regions

To investigate the possible AADk-substrate mimicry, we extensively studied the features of the pS/TQs in the substrates of the AADks.

In order to become phosphorylated, phosphorylation sites have to be located in the accessible sequence regions. This accessibility is usually accomplished by the integration of the site into an intrinsically disordered region of the protein, similarly to the examples shown in Figure 1. The majority of the experimentally verified pS/TQs are located in predicted disordered regions. Approximately 80% of the experimentally verified pS/TQs have an IUPred score ≥ 0.4 (Figure S9). It is worth noting that some of the pS/TQs (20%) which are in ordered regions might project from the accessible region of a secondary structure element. However, as there is insufficient structural data to back up this hypothesis, we focused our analysis to the S/TQs located in disordered regions.

The analysis of the sequences of the AADk substrates revealed that S/TQs in the disordered regions of the AADk substrates are more conserved than the S/TQs in proteins that do not interact with the AADks (Table S2). A similar trend was found for QS/Ts (Table S2). Interestingly, for TQ and QT we did not find a significant difference (Table S2). We also noticed that in the sequence XXXXXXXpS/TQXXXXXX, where pS/T is the experimentally validated phosphorylated residue, 60.54% of the cases have at least one additional verified phosphorylation site. The number of these additional sites varies from 1 (47.51%) to 6 (1%); furthermore, they are equally distributed N- or C-terminally to the pS/T.

Should the effectors be phosphorylated by the AADks, they can be expected to display these characteristics.

### Bacterial effectors might mimic AADk cellular substrates

We noticed that bacterial effectors resemble AADk substrates in terms of motif distribution: conserved S/TQs often appear close to conserved QS/Ts in disordered protein regions (Figure 1, Figure 4, Figure S5-S7). Furthermore, these sites are embedded in sequence regions that generally resemble host proteins. To show this, we checked for similarities between the amino acid compositions of the disordered regions of human proteins and selected pathogenic bacterial proteomes (*Legionella pneumophila, Chlamydia trachomatis, Coxiella burnetii* and *Pseudomonas syringae pv. tomato*). We did not find significant differences (Kruskal-Wallis p-value > 0.6 for all comparisons, Table S3), suggesting that while in general human and bacterial sequence compositions can highly differ, this difference is minimal considering disordered protein regions. It should be noted that bacteria use far fewer disordered proteins, but those they do, appear to have similar characteristics to the human disorderome.

We identified 10 bacterial effectors from human and plant pathogens that showed properties similar to the AADk multi-site substrates (Table S1; Figure 3D). We found eight different effectors from human pathogens: yhhA and Tir from enteropathogenic *Escherichia coli;* map from *Escherichia coli* O103:H2 (Verotoxigenic *E. coli);* CagA from *Helicobacter pylori*; sidG, lpg2844 and lpg2577 from *Legionella pneumophila*; and VPA1357 from *Vibrio parahaemolyticus*. yhhA is an experimentally validated effector from EPEC (47) but its function remains unknown. Interestingly, the sequence alignment generated using yhhA and its homologs shows that ST/Q and QS/T sites are well conserved across different pathogenic species (Figure 4A). We also noticed that in several cases, S/TQ and QS/T sites are conserved among closely related pathogens, and they even persist in the more divergent species, although their locations in the sequence may change (Figure S5-S7). Among the plant effectors, PopA and PopB are two experimentally validated cases (48,49) from *Ralstonia solanacearum*, an economically important pathogen that infects several crops. PopB also shows conserved ST/Q and QS/T motifs across different members of *Ralstonia* genus (Figure 4B).

Taking all these data into consideration, many bacterial effectors appear to have the properties of eukaryotic AADk substrates and may be mimicking their cellular substrate S/TQ motifs.

## DISCUSSION

Bacteria can mimic features of eukaryotic proteins in order to take advantage of the molecular machinery of the host cell during infection. This concept is well established for effectors such as Tir from the Enteropathogenic *Escherichia coli* (EPEC) (35). The C-terminal region of Tir shares similarity with the cellular Immunoreceptor Tyrosine-based Inhibition Motifs (ITIMs) which play a pivotal role in actin polymerisation. We have now found that bacterial effectors show conserved S/TQ motifs across different species of pathogenic bacteria (Figure 4), marking them as potential substrates of human AADks. During our analysis we observed that conserved S/TQs are present in both human (*Legionella pneumophila*, *Escherichia coli*, *Vibrio parahaemolyticus* and *Helicobacter pylori*) and plant pathogens *(Ralstonia solanacearum)*. Given that S/TQs are phosphorylated by AADks, we suppose that these sites could be phosphorylated by the host kinases. There is evidence that some bacteria, such as *Salmonella typhi* (50) and different strains of *E.coli* possess toxins with dsDNase activity such as the CDT toxin (51). Moreover, some plant pathogens have been reported to be able to cause DNA double-strand breaks, and ATM and ATR double mutants are more susceptible to infection than the wild-type plants (52). However, many more effectors with SQ motifs are present in human and plant pathogens that may not be directly damaging the host cell DNA.

For example, we found that *Shigella dysenteriae*, Enterohemorrhagic *E. coli* and EPEC all possess a small S/TQ-rich effector (yhhA and map) which might interact with ATM, interfering with its signalling pathway, resulting in a possible advantage for the bacteria. Oxidative stress is an essential factor for gastrointestinal mucosal diseases caused by pathogenic bacteria such as *Shigella* or *E.coli* (53). ATM plays a key role in oxidative stress response (4; 5) and the activation of its downstream pathway could either compromise, or enhance, the entire infection process. This hypothesis is supported by the cytoplasmic presence of AADks and their possible involvement in immune response (10-14) suggesting that their function is not limited to DNA damage response or cell cycle regulation. While the majority of the phosphorylation sites in eukaryotes must be accessible to the kinase (Figure S9) being in disordered regions or on the accessible surface region of domains, in bacteria some of the SQ sites could be phosphorylated even though they are embedded in ordered regions due to the fact that type III effectors enter unfolded into the target cell (54). Nevertheless, in order to obtain strong candidates, we decided to only consider candidate motifs in disordered regions during our analysis. Our analysis was complicated by the observation that QS/T motifs seemed to be associated with the S/TQ sites. We were able to show that these two motifs do significantly cluster together in both the cellular substrate sets and in the bacterial effectors. We were also quickly able to find multiple examples of S/TQ-rich proteins in both animal- and plant-infecting pathogens.

Most of these proteins are not yet known to be effectors (data not shown) though. The alignments show that both in human cellular proteins and in the bacterial proteins, when there are S/TQ clusters, the QS/T motif always clusters too. The function for QS/T is still unknown but the clustering suggests it could have a function linked to the phosphorylation of S/TQ. We noticed, in addition, that the theoretical distribution of S/TQ – QS/T distances differ significantly from the observed one. An experimentally validated docking motif for PIKK kinases was reported by Falk et al. (2005) but is only present in a few of the known nuclear substrates (10 out of 152). QS/T seems too simple to be a pure docking motif, such as the one found in MAPK substrates (55).

One possible role could be to act as a docking-activating or (perhaps less likely) a docking-inhibitory site. However, a few of the known cellular substrates (12 out of 152) do not have any QS/Ts in their sequences. This includes the key DNA damage protein H2AX. In such a scenario, the motif might work *in trans* because, before it can phosphorylate H2AX, ATM is docked to the damage site by NBS1 (56), which is itself also an ATM substrate with 6 copies of the QS/T motif.

Given the somewhat weak statistical results we were worried about the obvious temptation to try too many different tests. This would inevitably lead to a multiple testing problem. Finally, the motifs in the bacterial proteins show a broad range of conservation, with some motifs being very conserved and others partially or poorly. But in many cases, motifs would persist in all the sequences even if they were not conserved. It is well known that SLiMs can change position by evolutionary resampling: duplication followed by loss of the original (57). This suggests that there is a need to develop an evolutionary “motif persistence” score for these very simple sequence motifs.

In the preprint, we have reported several pieces of evidence that could suggest an interplay between bacterial effectors and the ATM kinase family. Bioinformatics analyses do not prove the function of a motif candidate and, as a consequence, they must be experimentally validated. If the AADk usage proves to be widespread in pathogens, then the oxidative stress signaling is likely to be an important part of that function. It will need to be established whether the pathogens actively promote oxidative stress to their advantage (53) or whether it is a side effect of infection that needs to be controlled. Tissue culture experiments with oxidants like hydrogen peroxide or antioxidants such as N-acetyl cysteine might show altered infectivity. In addition, the availability of ATM kinase inhibitors such as KU-55933 (58) will be useful for phosphorylation experiments.

## ACKNOWLEDGMENTS

The authors thank Dr. Bernd Klaus for discussing statistical issues. DS thanks fellowship from the Erasmus+ Program. BM acknowledges support from the EMBO|EuropaBio fellowship 7544. MK is thankful to the Humboldt Foundation for post-doctoral research fellowship.

